# Why slow axonal transport is bidirectional – can axonal transport of tau protein rely only on motor-driven anterograde transport?

**DOI:** 10.1101/2022.07.31.502201

**Authors:** Ivan A. Kuznetsov, Andrey V. Kuznetsov

## Abstract

Slow axonal transport (SAT) moves multiple proteins from the soma, where they are synthesized, to the axon terminal. Due to the great lengths of axons, SAT almost exclusively relies on active transport, which is driven by molecular motors. The puzzling feature of slow axonal transport is its bidirectionality. Although the net direction of SAT is anterograde, from the soma to the terminal, experiments show that it also contains the retrograde component. One of the proteins transported by SAT is microtubule-associated protein tau. To better understand why the retrograde component in tau transport is needed, we used the perturbation technique. We analyzed the simplification of the full tau SAT model for the case when retrograde motor-driven transport and diffusion-driven transport of tau are negligible, and tau is driven only by anterograde (kinesin) motors. The solution of the simplified equations shows that the tau concentration along the axon length stays almost uniform (decreases very slightly), which is inconsistent with the tau concentration at the outlet boundary (at the axon tip). Thus kinesin-driven transport alone is not enough to explain the experimentally observed distribution of tau, and the retrograde motor-driven component in SAT is needed.

## 1. Introduction

Due to the great length of axons, neurons have to rely on a complicated “railway” system composed of microtubules (MTs). Various cargos are transported along MTs being pooled by molecular motors. Typically, newly synthesized cargos are transported in the anterograde direction (from the soma to the axon terminal) by molecular motors that belong to the kinesin superfamily. Cargos that need to be recycled in the somatic lysosomes are transported in the retrograde direction by molecular motors that belong to the dynein superfamily. Axonal transport is classified into fast anterograde axonal transport, which is driven by kinesin motors (representative velocity 1 μm s^-1^), fast retrograde axonal transport, which is driven by dynein motors (representative velocity is also 1 μm s^-1^), and slow axonal transport (SAT) (Morfini et al. 2012; Brown, Anthony 2016; Roy 2020; Sleigh et al. 2019; Guedes-Dias and Holzbaur 2019).

One intriguing feature of SAT is its slow velocity. Cytoskeletal elements, such as neurofilaments (NFs), which are transported in slow component-a (SCa), move with an average anterograde velocity of 0.002– 0.02 μm s^-1^. Many other cargos, such as cytosolic proteins and tau protein (hereafter tau), which are transported in slow component-b (SCb), move with an average anterograde velocity of 0.02–0.09 μm s^-1^ (Brown 2000). It is puzzling because the average velocity of anterograde and retrograde molecular motors, kinesin and dynein, is approximately 1 μm s^-1^ (Brown 2000; Goldstein 2001). This puzzle was resolved independently in Wang et al. (2000) and Roy et al. (2000) who studied SCa transport of NFs and found that these cargos alternate between fast anterograde and retrograde movements (occurring with a velocity of ∼1 μm s^-1^) and frequent pauses. A representative portion of time that NFs spend pausing is 97% (Brown 2014), which explains the small average velocity of NFs.

The second mystery of SAT is that it moves cargoes in both anterograde and retrograde directions, although it has a net anterograde bias (Brown 2014). Since moving cargoes by molecular motors requires energy, the presence of the retrograde component in SAT is puzzling since it makes cargo delivery more energetically expensive. In this paper, we used mathematical modeling to explain why slow axonal transport must be bidirectional to allow for maintenance of concentration gradients along the axon length.

Tau is mostly known for its involvement in Alzheimer’s disease and other tauopathies when it becomes prone to aggregation (Huang et al. 2016; Iqbal et al. 2010; Zempell and Mandelkow 2014; Blennow et al. 2006). In axons, tau is transported by SCb (Utton et al. 2002; Utton et al. 2005; Brown, A. 2014). In this paper, we conducted the analysis on equations in a model developed in Kuznetsov and Kuznetsov (2017a, 2018, 2020) to investigate what happens when all other modes of transport except kinesin-driven transport of tau become negligible. Our goal was to continue research reported in Kuznetsov and Kuznetsov (2022, 2023a) for α-synuclein, as well as Kuznetsov and Kuznetsov (2023b) for MAP1B, and to determine whether anterograde motor-driven transport alone can replicate the tau distribution observed in experiments reported by Black et al. (1996). Focusing on scenarios where the dynein velocity and tau diffusivity were both small, we were able to derive an analytical solution for the tau transport model through the application of the perturbation technique. This solution can be useful for verifying the accuracy of numerical codes.

Since in a healthy neuron tau predominantly exists in the monomeric form, hereafter we denoted tau monomer as tau.

## 2. Materials and models

### 2.1. Mathematical model of tau transport in an axon

A schematic diagram of the axon is displayed in Fig. 1. Since we need equations describing tau transport for the perturbation analysis, we briefly restate here the equations developed in Kuznetsov and Kuznetsov (2017a, 2018, 2020).

**Fig. 1.**
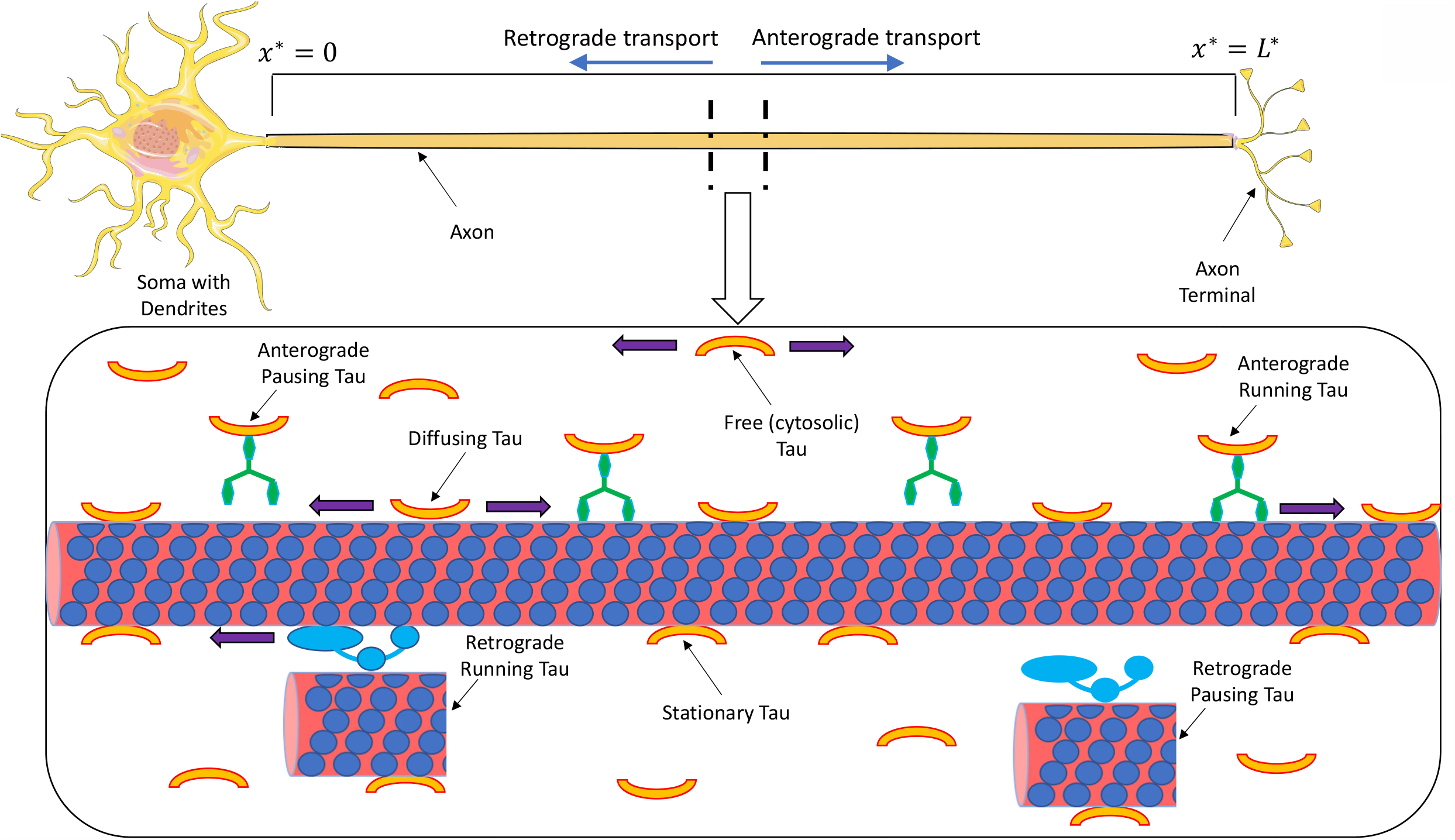
A diagram of a neuron showing various modes of SAT and diffusion of tau. In displaying possible mechanisms of tau transport, we followed Scholz and Mandelkow (2014) (see Fig. 3 in Scholz and Mandelkow 2014). Tau can be transported by molecular motors, kinesin and dynein (active transport), or by a diffusion-driven mechanism, either in the cytosol or by diffusion along MTs. Motor-driven retrograde transport of tau may occur by tau piggybacking on short microtubule fragments translocated by cytoplasmic dynein.

The model simulates tau transitions between seven different kinetic states (Fig. 2). A tau concentration in a corresponding kinetic state is given below in the parentheses. Two of the kinetic states describe active anterograde 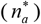 and retrograde 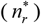 transport of tau propelled by kinesin and dynein motors, respectively (Utton et al. 2005; Butler et al. 2019). Two other kinetic states are anterograde 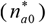 and retrograde 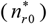 pausing states, respectively, when motors driving the cargo temporarily disengage from MTs, but are ready to re-engage and resume their motion. There are also two kinetic states in which tau can be driven by diffusion. These are free (cytosolic) state 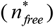 (Samsonov et al. 2004; Konzack et al. 2007; Weissmann et al. 2009) and a kinetic state representing a sub-population of MT-bound tau that can diffuse along the MTs 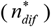 (Hinrichs et al. 2012). There is also a sub-population of MT-bound tau that is stationary 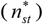 (Hinrichs et al. 2012). Since an axon is much longer in one dimension than in the other two dimensions, all tau concentrations in the axon are represented by one-dimensional densities of tau monomers (μm^-1^).

**Fig. 2.**
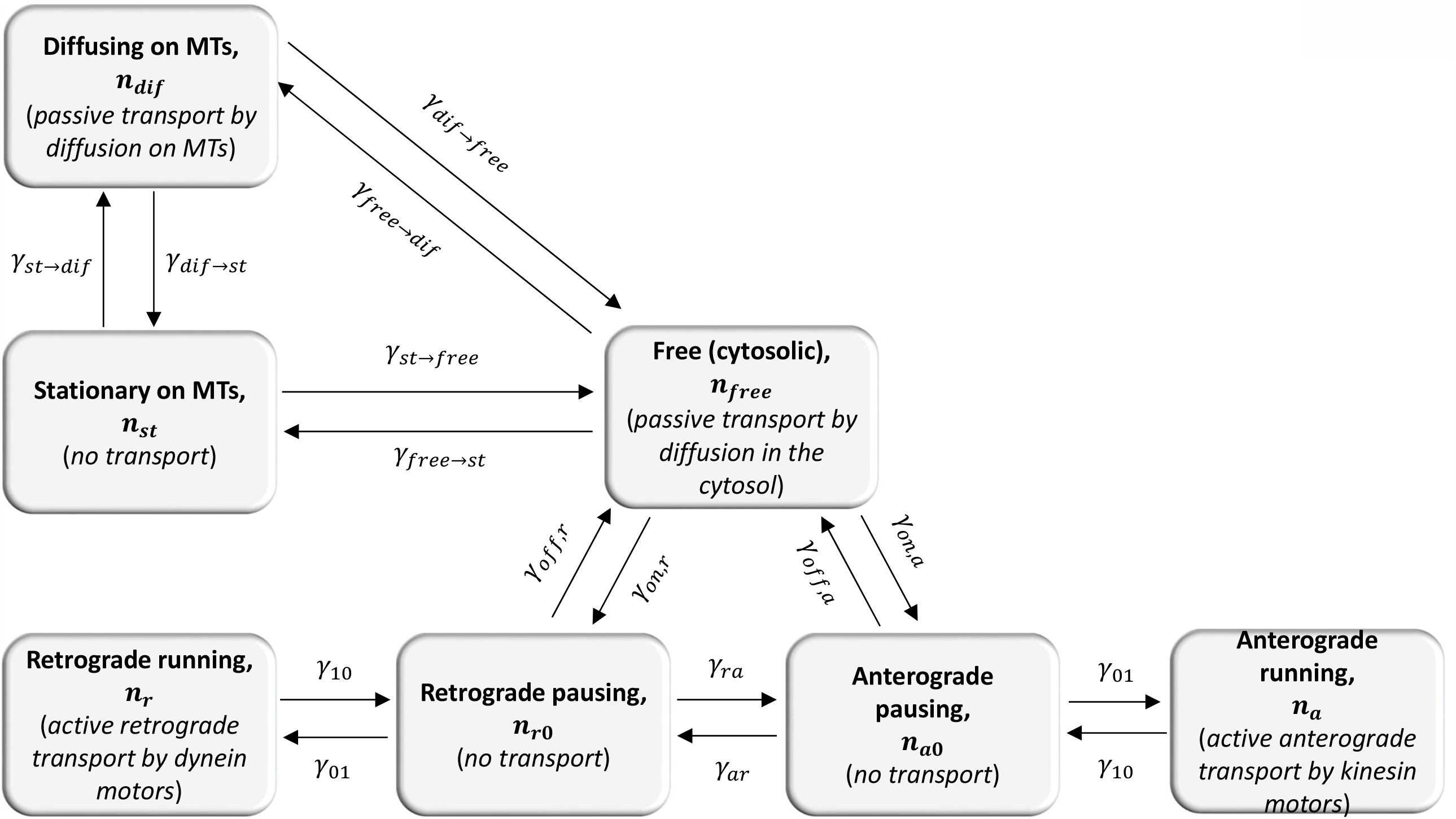
A kinetic diagram displaying seven kinetic states in our model of tau transport in the axon.

Tau interaction with MTs is highly dynamic. Single-molecule tracking experiments conducted by Janning et al. (2014) reveal that tau remains on a single microtubule for around 40 ms before hopping on to the next one. Since the diffusion timescale of tau is considerably larger than that of these hops, our model predicts time-averaged values of tau concentrations. This is similar to continuum theory, which does track the movement of individual molecules.

A Cartesian coordinate directed along the axon is denoted as *x*\* (Fig. 1). We used the SAT model of tau developed in Kuznetsov and Kuznetsov (2017a, 2018, 2020). The model consists of seven equations obtained by stating tau conservation in the corresponding kinetic states. We assume that in a healthy axon transport processes are quasi-steady; thus ∂… / ∂*t*\* = 0. The conservation of tau in anterograde and retrograde motor-driven states leads to the following equations, respectively (Fig. 2):

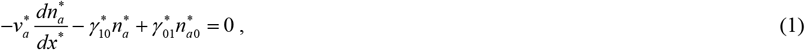

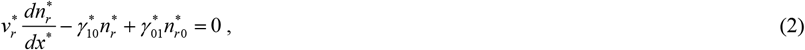

where asterisks denote dimensional quantities. The first terms on the left-hand sides of Eqs. (1) and (2) represent the change of tau concentration in a control volume due to motor-driven transport, while the terms that are multiplied by kinetic constants (*γ* * *s*) represent transitions of tau between motor-driven and pausing kinetic states (Fig. 2). We defined model parameters in Tables S1 and S2 in the Supplemental Materials.

Stating the conservation of tau in the pausing states, which simulate the situation when tau is temporarily disengaged from MTs, gives the following equations (Fig. 2):

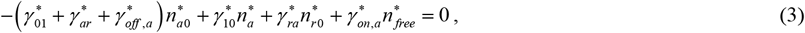

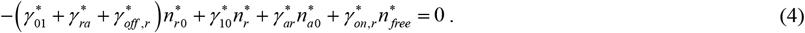

Because tau does not move in the pausing states, tau concentrations in the pausing states can change only due to tau transitions to and from other kinetic states.

Stating the conservation of tau in the free state, which simulates the pool of tau that is freely suspended in the cytosol and is not associated with MTs, gives the following equation:

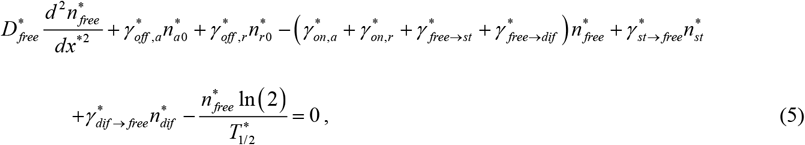

where the first term on the left-hand side of Eq. (5) simulates diffusion-driven transport of tau in the cytosol, and the last term on the left-hand side of Eq. (5) simulates tau degradation in proteasomes (Poppek et al. 2006; Kierszenbaum 2000). The tau degradation term is only included for the free (cytosolic) tau because tau must be detached from MTs to enter a proteasome’s proteolytic chamber. All other terms in Eq. (5) simulate tau transitions to/from the free cytosolic state.

It is important to mention that the model considers two possible mechanisms of tau protein transition between a retrograde pausing state (bound to dynein) and an anterograde pausing state (bound to kinesin), see Fig. 2. (i) Tau protein can first unbind from a kinesin and then bind to a dynein. In this case, the transition is through the free state. (ii) Tau protein bound to kinesin can be directly “stolen” by a dynein (and bind to this dynein), see Eqs. (3) and (4).

Tau that is attached to MTs can be divided into stationary tau and tau that can diffuse along the MTs (Hinrichs et al. 2012). Stating the conservation of the diffusing fraction of MT-bound tau gives the following equation:

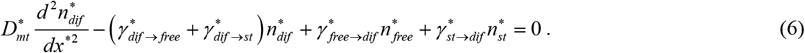

The first term on the left-hand side of Eq. (6) simulates transport of tau by its diffusion along MTs. All other terms simulate transitions to/from the kinetic state representing tau diffusing along MTs (Fig. 2).

Part of tau attached to MTs is stationary (Hinrichs et al. 2012), and stating tau conservation in the stationary kinetic state leads to the following equation:

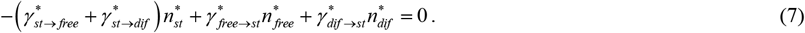

Since tau in the stationary state does not move, its concentration can only change due to tau transitions to/from the stationary state (Fig. 2).

The total tau concentration is calculated by finding the sum of tau concentrations in all seven kinetic states (Fig. 2):

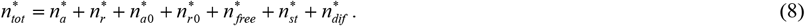

Tau is associated with MTs in six out of seven kinetic states (except for 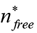). The percentage of MT-bound tau is found by calculating the ratio of the sum of tau concentrations in six MT-associated states to 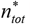:

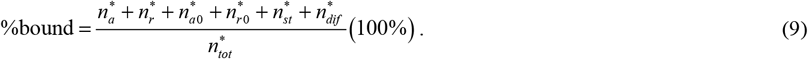

Tau can be driven by four mechanisms: anterograde transport by kinesin motors, retrograde transport by dynein motors, diffusion of free tau in the cytosol, and diffusion of MT-bound tau along MTs. The total flux of tau can be found as:

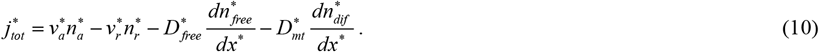

The average tau velocity was found as suggested in Kuznetsov and Kuznetsov (2015):

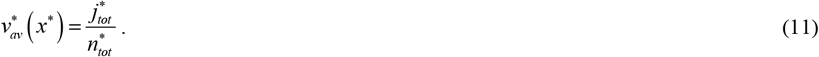

Eqs. (1)-(7) require six boundary conditions. We imposed three boundary conditions at the axon hillock:

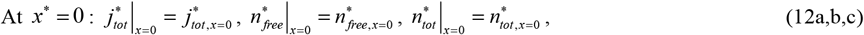

which postulate the tau flux into the axon, the concentration of free tau at the axon hillock, and the total concentration of tau at the axon hillock, respectively. *j*_*tot, x*=0_ and *n*_*free, x*=0_ are determined by finding values that give the best fit between model predictions and published experimental results (Table S2). The following three boundary conditions were imposed at the axon terminal:

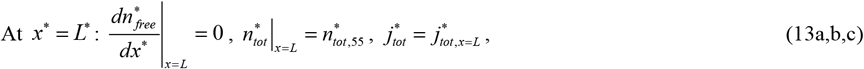

which postulate zero concentration gradient of free tau at the axon tip, the total tau concentration at the axon tip, and the flux of tau into the axon terminal, respectively. The flux of tau into the axon terminal, 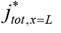, in Eq. (13c) was calculated according to Eq. (S1), see section S2 in the Supplemental Materials.

Some of the values of the model parameters can be found in the literature; these are summarized in Table S1. Other parameters whose values are not found in the literature (for example kinetic constants characterizing the rates of tau transitions between different kinetic states, *γ* * *s*) are summarized in Table S2. These parameters in Table S2 were determined by finding values that give the best fit with published experimental data. Our problem involves different types of published data, for example, the tau concentration and velocity. Therefore, we used multi-objective optimization (Kool et al. 1987; Zadeh 2008; Zadeh and Shah 2010; Zadeh 2011; Zadeh and Montas 2014) to determine best-fit values of these parameters in Table S2. The following objective (penalty) function, which characterizes the deviation of model predictions from published data, was utilized:

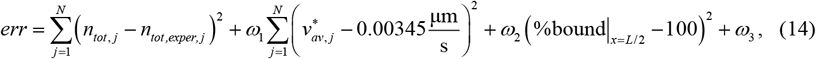

where the dimensionless total tau concentration as well as other dimensionless parameters are defined in Table S3. Eqs. (1)-(7) with boundary conditions (12), (13) were solved for various values of parameters given in Table S2. This was done until the values of parameters that give the global minimum to the objective function defined in Eq. (14) were found. More details about the minimization algorithm are given in section S3.

The first term on the right-hand side of Eq. (14) characterizes the deviation between the predicted and experimentally measured tau concentrations along the axon length. The experimentally measured concentration was obtained by scanning the tau concentration data reported in Fig. 7D of Black et al. (1996) in 55 (*N*=55) experimental points.

The second term on the right-hand side of Eq. (14) characterizes the deviation between the predicted velocity of tau transport (see Eq. (11)) and the average of the reported range of tau velocity (0.2-0.4 mm/day, Mercken et al. (1995), Scholz and Mandelkow (2014)).

The third term on the right-hand side of Eq. (14) is utilized to simulate that most of tau must be attached to MTs. (According to Janning et al. (2014), approximately 99% of tau is bound to MTs.) We set *ω*_1_ =10,000 s^2^/μm^2^ and *ω*_2_ =1. This was done based on extensive numerical trials to avoid overfitting the tau concentration, average velocity, or the percentage of MT-bound tau. Note that due to dimensions of *ω*_1_ the second term on the right-hand side of Eq. (14) is dimensionless.

The fourth term on the right-hand side of Eq. (14) prevents tau concentrations in any of the kinetics states displayed in Fig. 2 from becoming negative, which would be unphysical. The non-negative values were achieved by setting *ω*_3_ to a large positive value, 10^8^, if any of *n*_*a,j*_, *n*_*r,j*_, *n*_*a*0,*j*_, *n*_*r*0,*j*_, *n*_*free,j*_, *n*_*diff,j*_, or *n*_*st, j*_ at any location in the axon (*j*=1,…,55) becomes negative. Otherwise, *ω*_3_ was set to zero. This allowed the reformulation of a constrained optimization problem with a non-negative constraint into an unconstrained optimization problem with a penalty function (Yang and Huang 2004).

Numerical solution procedure of Eqs. (1)-(7) is outlined in section S3. The method of finding the best-fit values of parameters given in Table S2 by minimizing the penalty function given by Eq. (14) is also described in section S3.

The full model of slow axonal transport of tau given by Eqs. (1)-(7), (12), (13), and (S1) contains 17 parameters whose values must be determined by minimizing the discrepancy between model predictions and published experimental data. The number of model parameters can be reduced by identifying parameters to which the numerical solution is least sensitive Kuznetsov and Kuznetsov (2019a, 2019b). A similar approach was used in Kuznetsov and Kuznetsov (2017a) to obtain a reduced form of the model that contains only 8 fit parameters. Nevertheless, for the purposes of the research presented in this paper, the reduced model is not necessary since the full model cannot simulate the increase of tau concentration towards the axon tip without retrograde motors, as we are demonstrating here. If the complete and more comprehensive model is incapable of achieving this, the reduced model will also not be able to accomplish this task either.

### 2.2 A perturbation solution of the tau transport model for the case of small dynein velocity and small tau diffusivity

We are interested in investigating whether the model can simulate cargo transport against the cargo’s concentration gradient by kinesin-driven transport alone. This requires setting the dynein velocity and the tau diffusivity, both in the cytosolic state and in the state that represents tau that diffuses along MTs, to small values. For this purpose, the following small parameters were introduced. The dimensionless retrograde velocity was defined as:

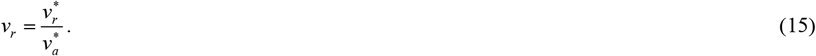

The dimensionless cargo diffusivity in the cytosolic state is defined as:

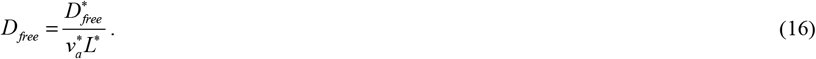

The right-hand side of Eq. (16) represents the reciprocal to the Peclet number. Eq. (16) can be re-written as:

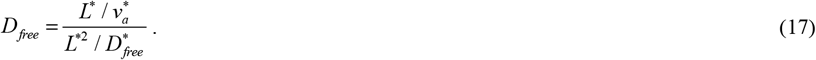

Thus, *D*_*free*_ can be viewed as the ratio of the convection time to diffusion time for free tau.

Similarly, the dimensionless cargo diffusivity of MT-bound tau that diffuses alone MTs is defined as:

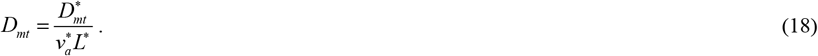

Since

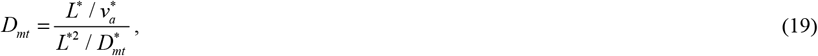

*D*_*mt*_ can be interpreted as the ratio of the convection time to diffusion time of MT-diffusing tau.

If the dynein velocity is small, *v*_*r*_ can be treated as a small parameter. For seven dependent variables of the model, 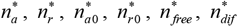, and 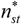, we used the following perturbation expansions (for example):

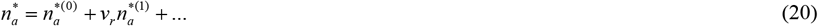

We then substituted the perturbation expansions (such as Eq. (20)) for all seven dependent variables into Eqs. (1)-(7). We separated the terms that do and do not contain the small parameter *v*_*r*_ in the obtained equations and equated the terms that do not contain *v*_*r*_ to zero (the terms containing *v*_*r*_ were neglected). As a result, Eqs. (S2)-(S8) were obtained.

If the diffusivity of cytosolic tau is small, *D*_*free*_ can be treated as a small parameter. For seven dependent variables of the model, 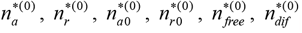, and 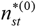, we used the following perturbation expansions (for example):

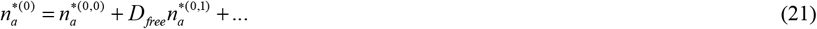

We then substituted the perturbation expansions (such as Eq. (21)) for all seven dependent variables into Eqs. (S2)-(S8). We separated the terms that do and do not contain the small parameter *D*_*free*_ in the obtained equations, and equated the terms that do not contain *D*_*free*_ to zero (the terms containing *D*_*free*_ were neglected). As a result, Eqs. (S9)-(S15) were obtained.

If the diffusivity of MT-bound tau is small, *D*_*mt*_ can be treated as a small parameter. For seven dependent variables of the model, 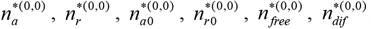, and 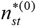, we used the following perturbation expansions (for example):

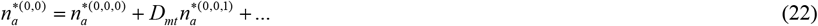

We then substituted the perturbation expansions (such as Eq. (22)) for all seven dependent variables into Eqs. (S9)-(S15). We separated the terms that do and do not contain the small parameter *D*_*mt*_ in the obtained equations and equated the terms that do not contain *D*_*mt*_ to zero (the terms containing *D*_*mt*_ were neglected). As a result, Eqs. (S16)-(S22) were obtained.

To simplify the notations, we replaced the superscript (0,0,0) with [0]. By solving Eq. (S17) for 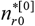, the following equation was obtained:

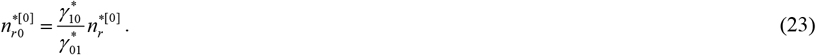

We used Eq. (23) to eliminate 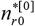 from Eqs. (S16), (S18)-(S22), and obtained Eqs. (S23)-(S28).

By eliminating 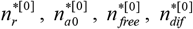, and 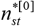 from Eqs. (S23)-(S28), we obtained the following equation for 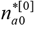:

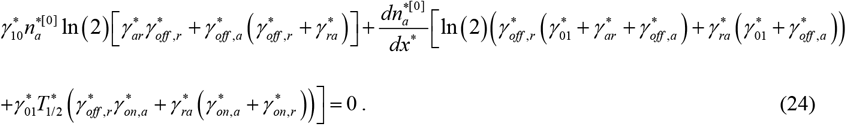

To compare the numerical solution of Eqs. (1)-(7) and the analytical solution of the perturbation equations (S15)-(S21) we have to be selective about the boundary condition. While Eqs. (1)-(7) require six boundary conditions, which are postulated by Eqs. (12) and (13), the perturbation equations (S15)-(S21) allow for only one boundary condition. To compare the two solutions, we assumed the same rate of tau synthesis in the soma. This results in the same tau flux from the soma into the axon, 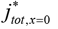. We thus solved Eq. (24) subject to the following boundary condition, which is identical to Eq. (12a):

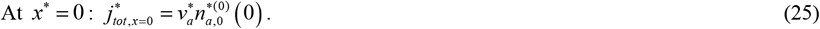

Eq. (25) is valid because in the kinesin-only transport model, the flux of tau is calculated as:

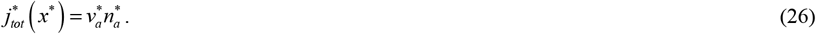

The solution of Eq. (24) subject to boundary condition (25) is

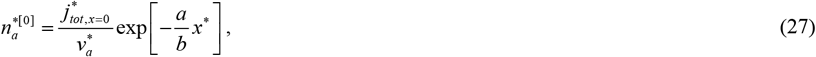

where

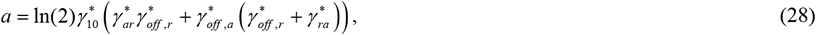

and

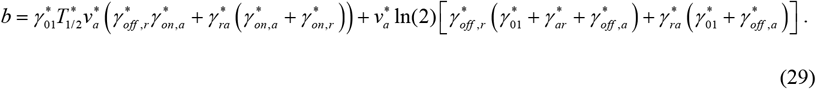

Solving Eq. (S23) for 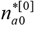, the following is obtained:

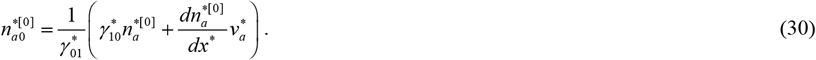

By eliminating 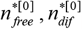, and 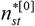 from Eqs. (S24), (S25), (S27), and (S28), the following equation for 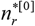 is obtained:

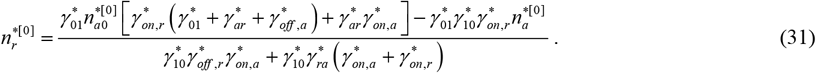

It should be noted that although the perturbation solution is obtained for *v*_*r*_ → 0, the tau concentration 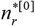 is still defined, tau just does not move into this kinetic state. The same applies to tau concentrations in the diffusion-driven states, 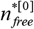 and 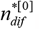.

By eliminating 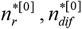, and 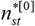 from Eqs. (S24), (S25), (S27), and (S28), the following equation for 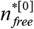 is obtained:

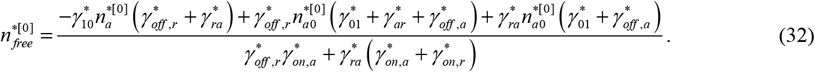

By eliminating 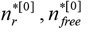, and 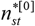 from Eqs. (S24), (S25), (S27), and (S28), the following equation for 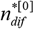 is obtained:

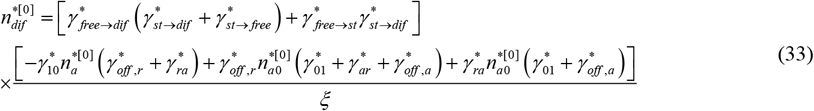

where

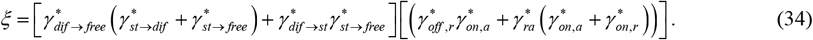

By eliminating 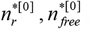, and 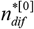 from Eqs. (S24), (S25), (S27), and (S28), the following equation for

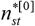 is obtained:

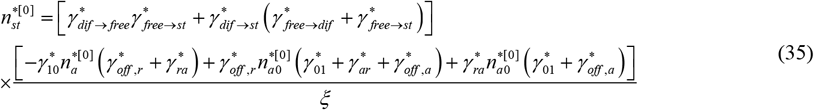

## 3. Results

In all numerically obtained figures in this paper (all figures except the definition sketches given in Figs. 1 and 2), we used the same values for parameters given in Table S1 to compute the curves that represent the full SAT model and the model that accounts for anterograde (kinesin-driven) transport only. This is done to enable a comparison between predictions of the two models. The values of the parameters in Table S2 are computed as values that minimize the penalty function given by Eq. (14). Note a good agreement between the model predictions and experimental results, which suggests that the optimization procedure described in section S3 worked correctly. The same values of these parameters are then used for computing the curves representing the anterograde-only transport model. This is done to ensure that the physical consequences of removing the dynein velocity and tau diffusivity from the model are not masked by the curve-fitting procedure. The curve-fitting procedure attempts to adjust the parameter values to minimize the discrepancy between the results of the model and the experimental results. An attempt to curve-fit the parameters every time the value of dynein velocity (for example) is changed would result in simultaneous change of all 18 parameters (dynein velocity plus 17 adjustable constants), and it would be difficult to determine what is causing the change in tau distribution.

All tau concentrations were non-dimensionalized by dividing them by the total tau concentration at the axon hillock (at *x*\* = 0), as shown in Table S3. The full SAT model can simulate experimentally predicted variation of the total tau concentration perfectly well. However, a model that accounts only for kinesin-driven transport predicts a uniform distribution of tau along the axon length (Fig. 3a), which suggests that an additional tau transport mechanism is needed to simulate a variation of tau concentration along the axon length.

**Fig. 3.**
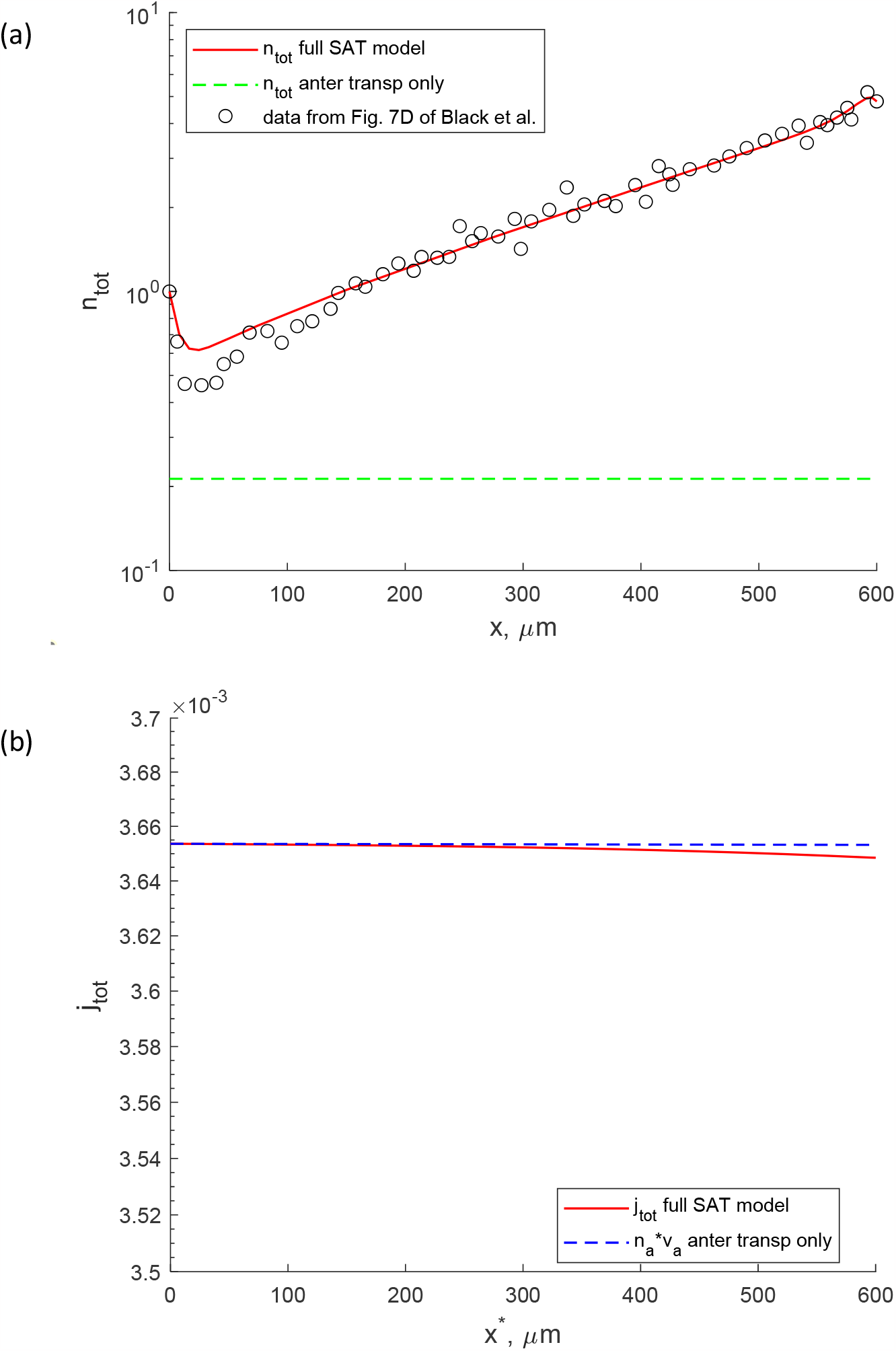
(a) Total concentration of tau. The solution for a full SAT model, which accounts for four mechanisms of tau transport (anterograde and retrograde SAT as well as tau diffusion in the cytosol and along MTs), was obtained numerically. The solution for a model that accounts for anterograde (kinesin-driven) transport only was obtained analytically, since for this case the closed-form solution given by Eqs. (23), (27), and (30)-(34) is available. Experimental data from Fig. 7D of Black et al. (1996) are shown by open circles. These data were rescaled such that the experimentally measured tau concentration at the axon hillock (at *x*\* = 0) was equal to unity. (b) Total flux of tau versus position in the axon, for the full SAT model and the model that simulates anterograde motor-driven transport only.

Both the full SAT model and kinesin-only driven model assume the same flux of tau entering the axon from the soma (see Eqs. (12a) and (25)). This explains why both curves in Fig. 3b cross at *x*\* = 0. A little faster decay of the curve depicting the tau flux predicted by the full SAT model (Fig. 3b) occurs because tau can be degraded in proteasomes only in the free (cytosolic) state, and the full SAT model predicts a larger concentration of tau in the free state than that predicted by the anterograde-only model (Fig. 4a).

**Fig. 4.**
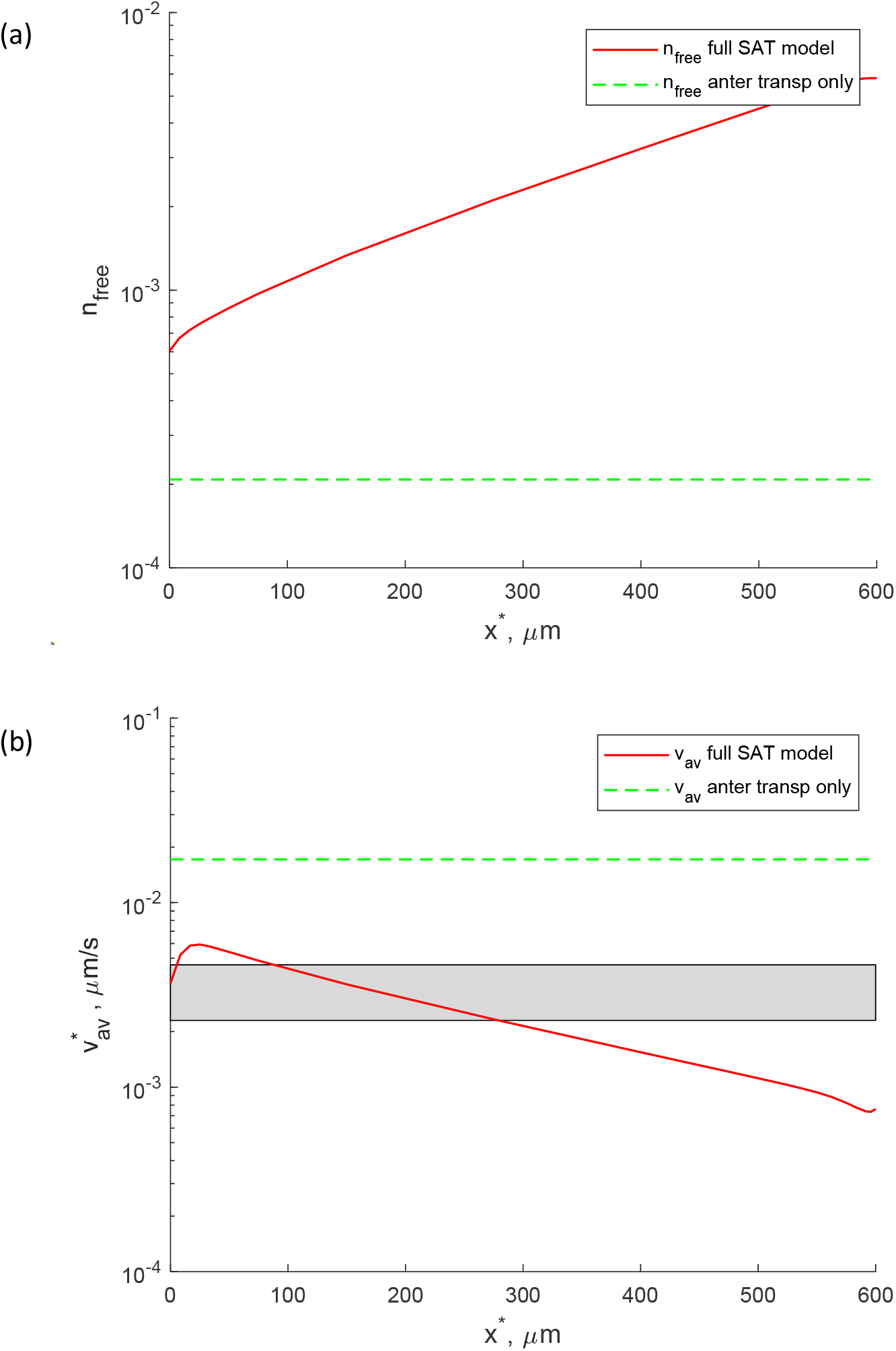
(a) Concentration of cytosolic tau that can freely diffuse and can be degraded in proteasomes. (b) Average velocity of tau. The range of the average tau velocity reported in Mercken et al. (1995) is shown by a horizontal band. Distributions for the full SAT model and the model that simulates anterograde motor-driven transport only are displayed.

According to the full SAT model, the following trends are predicted. The concentrations of anterograde and retrograde motor-driven tau stay constant over most of the axon, then sharply increase close to the axon terminal (Fig. S1). Distributions of anterograde and retrograde pausing tau behave similarly (Fig. S2). Concentrations of diffusing and stationary MT-bound tau first decrease, then linearly increase, and then decrease again. This suggests that tau diffusion works against anterograde kinesin-driven transport of tau over most of the axon (Fig. S3). If the model accounts only for anterograde kinesin-driven transport of tau, all the above tau concentrations are predicted to be uniform along the axon length (Figs. S1-S3).

The average velocity of tau transport is calculated as the ratio of the total flux of tau to the total tau concentration (Eq. (11)). The flux of tau decays very slowly due to a small rate of tau degradation in proteasomes (Fig. 3b). Therefore, the average velocity of tau (Fig. 4b) is inversely proportional to the total tau concentration displayed in Fig. 3a. This explains the shape of the tau average velocity for the full SAT model. For the anterograde-only transport model, the average tau velocity is given by a horizontal line (Fig. 4b).

The percentage of MT-bound tau, defined by Eq. (9), is close to 100% over the whole length of the axon (Fig. 5a). This is because the best-fit values of model parameters given in Table S2 are computed such that this criterion would be satisfied (see the third term of the penalty function defined by Eq. (14)).

**Fig. 5.**
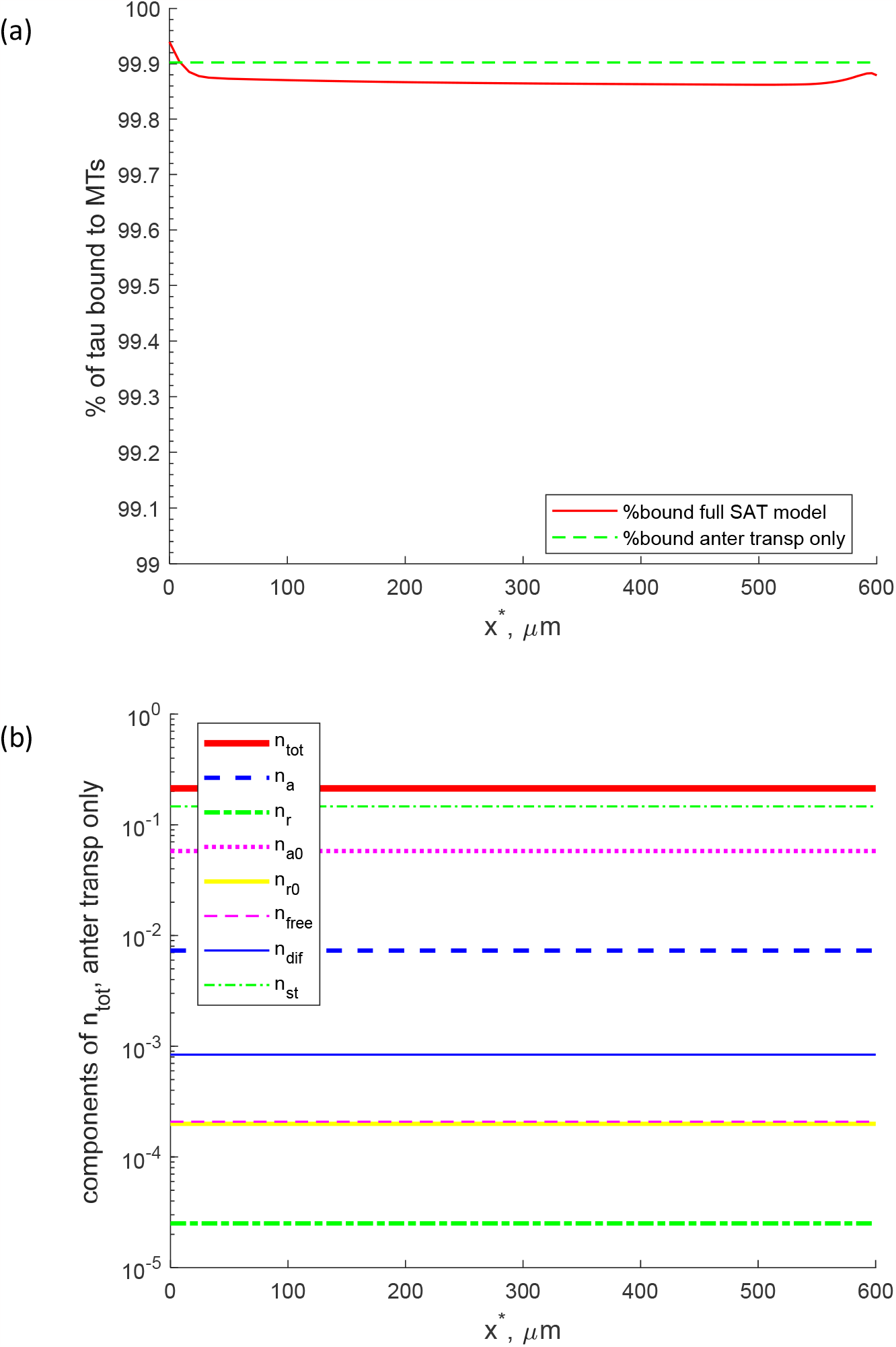
(a) Percentage of MT-bound tau versus position in the axon, for the full SAT model and the model that simulates anterograde motor-driven transport only. (b) Total concentration of tau and seven of its components for the model that simulates anterograde motor-driven transport only. The fact that the model with anterograde motor-driven transport only predicts tau distributions that are given by horizontal lines suggests that accounting for other mechanisms of tau transport (in addition to anterograde motor-driven transport) is required to explain a variation of tau distribution along the axon length.

In Fig. 5b we plotted the distributions of the total tau concentration and seven of its components predicted by the model that accounts only for kinesin-driven transport of tau. For the parameter values given in Tables S1 and S2, 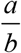 in Eq. (27) equals to 1.34 ×10^−9^ μm^-1^, which means that *n*_*a*_ is almost uniform, as shown in Fig. 5b. A slight decay of *n*_*a*_ is caused by a non-infinite half-life of tau. If the half-life of tau is increased to 2.16×10^10^ s (which is practically infinite), 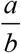 in Eq. (27) decreases to 1.37 ×10^−14^ μm^-1^. The same applies to tau concentrations in all other kinetic states. Since the total tau concentration is the sum of tau concentrations in all kinetic states, the total tau concentration is also uniformly distributed along the axon length (Fig. 5b). Hence, to simulate a variation of tau concentration along the axon length, another tau transport mechanism, in addition to kinesin-driven transport, is needed.

## 4. Discussion, limitations of the approach, and future directions

Experimental studies conducted by Wang et al. (2000) and Roy et al. (2000) have independently demonstrated that slow axonal transport includes both anterograde and retrograde transport, with long pauses in between motor-driven motions of cargo. However, why slow axonal transport has two components (anterograde and retrograde) remains a mystery. Our findings suggest that cargo distribution experimentally observed in slow axonal transport cannot be reproduced without retrograde transport or diffusion. Since the diffusivity of most of the slow axonal transport cargos during their axonal transport is small, this explains why retrograde transport is necessary for slow axonal transport.

To validate our hypothesis, we used the following approach. We utilized the full model of slow axonal transport (given by Eqs. (1)-(7)), numerically solved it, and optimized a penalty function to determine unknown parameters. We investigated which equations the full slow axonal transport model reduces to if retrograde motor-driven transport and tau diffusion become negligibly small. In this scenario, the only remaining mechanism for tau transport in the axon is kinesin-driven anterograde transport. Then we analytically solved Eqs. (S16)-(S22) of the simplified model with the inferred parameters. Finally, we compared the solutions of both models to published experimental measurements and determined that the simplified model without retrograde transport and diffusion was unable to adequately fit the experimental results. Indeed, the simplified model resulted in close to uniform tau concentrations in all kinetic states along the length of the axon (as shown in Fig. 5b). (The simplified model produced a concentration gradient that is exponential in *x*, but it is negligibly small given the fixed values of the parameters used in the model. Further, the tau concentration is strictly decreasing over the axonal domain, see Eq. (27), because of the degradation of unbound tau.) It became evident that a model that only considers anterograde motor-driven transport cannot simulate the prescribed tau variation along the axon length.

The reason is that Eqs. (S16)-(S22) contain only one ordinary differential equation, Eq. (S16). The rest of the equations are algebraic equations. The system is thus of the first order and allows only one boundary condition, which must be imposed at the axon hillock (at *x*\* = 0). As we suggested in Kuznetsov and Kuznetsov (2022, 2023a) (for α-synuclein) and in Kuznetsov and Kuznetsov (2023b) (for MAP1B), to be able to simulate a given cargo distribution in the axon, the model must allow for the imposition of the second boundary condition, which would allow to postulate a prescribed cargo concentration at the terminal (at *x*\* = *L*\*). Thus, a model that accounts for anterograde motor-driven transport alone cannot simulate a prescribed cargo variation in the axon. Non-monotonic distributions of tau at steady-state, such as those measured by Black et al. (1996), can only be obtained from a combination of transport mechanisms. For long-range transport, diffusion is not a good candidate for the second mechanism because it would take an extremely long time to transport a protein particle by a distance corresponding to the length of the axon. The effect of cargo diffusivity was studied in Kuznetsov and Kuznetsov (2015, 2017b, 2023a). It was established that for physiologically relevant values of cargo diffusivity the effect of diffusion-driven transport of cargo is primarily limited to two diffusion boundary layers. These layers are located at the ends of the axon, at the axon hillock and axon synapse.

The only remaining choice for the second mechanism is then retrograde dynein-driven transport. This conclusion has an important implication. It suggests why slow axonal transport is bidirectional, and why it contains both anterograde and retrograde motor-driven components. This is needed to support a cargo concentration that increases along the axon length, which requires cargo transport against its diffusion-driven flux.

Current research is limited because we used a continuum model for simulating cargo (tau) transport in the axon. Future research should involve simulating each cargo as an individual particle (Lai et al. (2018)). This would enable further testing of our hypothesis that anterograde-only motor-driven transport is insufficient for supporting cargo transport against its concentration gradient (when the cargo concentration at the tip of the axon is larger than at its hillock), and an additional transport mechanism, such as retrograde motor-driven cargo transport, is needed to support cargo transport in this situation. Possible effects of retrograde motor dysfunction on tau transport in Alzheimer’s disease (Kuznetsov and Kuznetsov 2018) also need to be investigated. The procedure of the determining values of the best-fit parameters given in Table S2 can also be improved in future research. One can envision performing a constrained optimization, where certain parameter values are required to stay within a defined range. These ranges can be obtained directly from existing literature or be set based on reasonable assumptions, such as being within 50% of the mean value.

## List of Abbreviations

LSR: least square regression
MT: microtubule
NF: neurofilament
SAT: slow axonal transport
SCa: slow component-a
SCb: slow component-b

## Acknowledgment

IAK acknowledges the fellowship support of the Paul and Daisy Soros Fellowship for New Americans and the NIH/National Institute of Mental Health (NIMH) Ruth L. Kirchstein NRSA (F30 MH122076-01). AVK acknowledges the support of the National Science Foundation (award CBET-2042834) and the Alexander von Humboldt Foundation through the Humboldt Research Award.

## Supplemental Materials

### S1. Supplementary tables

**Table S1.**
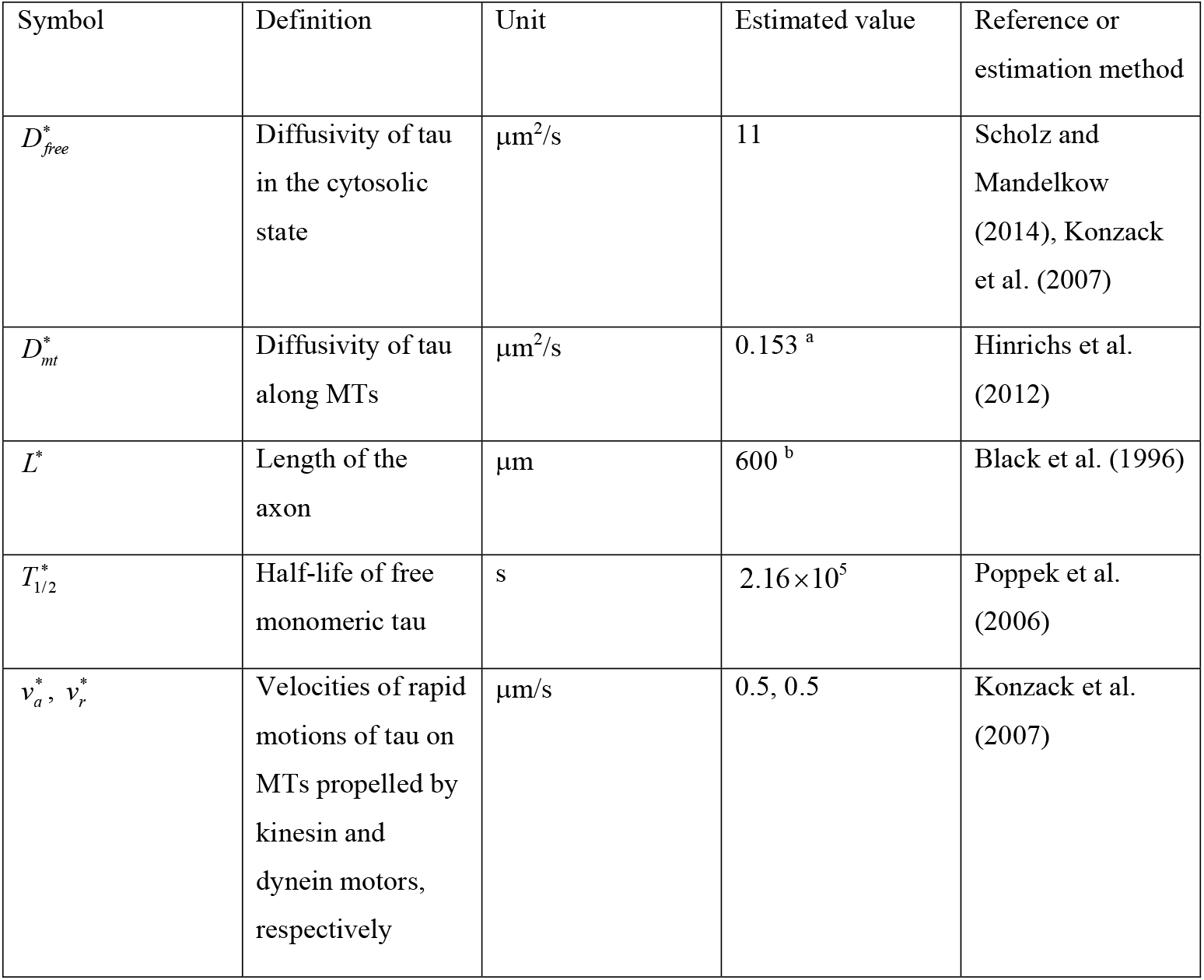
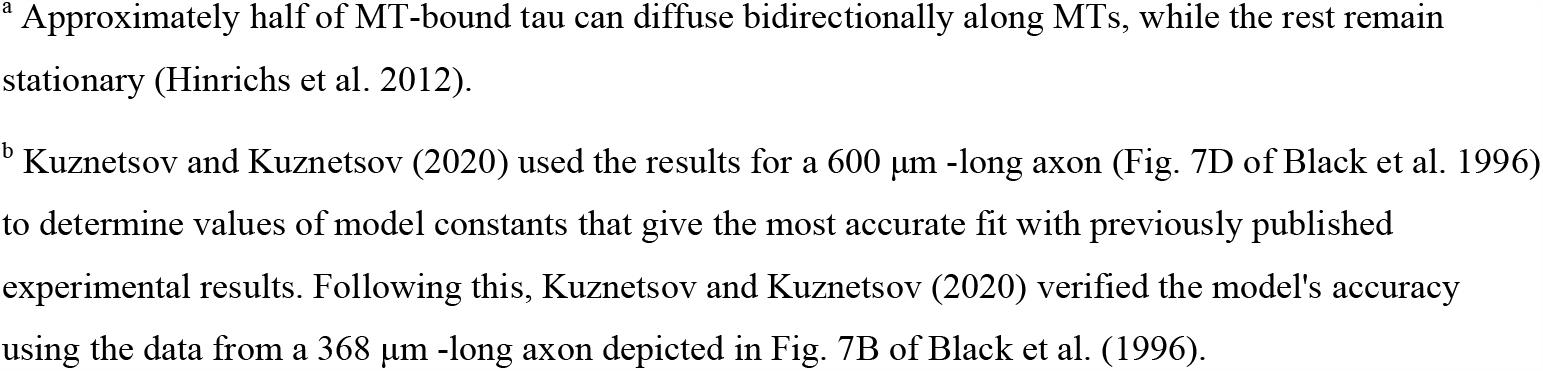
Parameters of the model that were estimated based on values found in literature.

| Symbol | Definition | Unit | Estimated value | Reference or estimation method |
| --- | --- | --- | --- | --- |
| $D_{free}^*$ | Diffusivity of tau in the cytosolic state | $\mu\text{m}^2/\text{s}$ | 11 | Scholz and Mandelkow (2014), Konzack et al. (2007) |
| $D_{mt}^*$ | Diffusivity of tau along MTs | $\mu\text{m}^2/\text{s}$ | 0.153 <sup>a</sup> | Hinrichs et al. (2012) |
| $L^*$ | Length of the axon | $\mu\text{m}$ | 600 <sup>b</sup> | Black et al. (1996) |
| $T_{1/2}^*$ | Half-life of free monomeric tau | s | $2.16 \times 10^5$ | Poppek et al. (2006) |
| $v_a^*, v_r^*$ | Velocities of rapid motions of tau on MTs propelled by kinesin and dynein motors, respectively | $\mu\text{m}/\text{s}$ | 0.5, 0.5 | Konzack et al. (2007) |

<sup>a</sup> Approximately half of MT-bound tau can diffuse bidirectionally along MTs, while the rest remain stationary (Hinrichs et al. 2012).
<sup>b</sup> Kuznetsov and Kuznetsov (2020) used the results for a 600 $\mu\text{m}$ -long axon (Fig. 7D of Black et al. 1996) to determine values of model constants that give the most accurate fit with previously published experimental results. Following this, Kuznetsov and Kuznetsov (2020) verified the model's accuracy using the data from a 368 $\mu\text{m}$ -long axon depicted in Fig. 7B of Black et al. (1996).

**Table S2.** Parameters that characterize transport of tau in the axon and their estimated values. The values reported here are different from those reported in Kuznetsov and Kuznetsov (2017a) because we now used a different value of 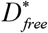. LSR (Least Square Regression), Beck and Arnold (1977).

| Symbol | Definition | Unit | Estimated value | Reference or estimation method |
| --- | --- | --- | --- | --- |
| $\gamma_{10}^*$ | Kinetic constant describing the rate of transitions $n_a^* \rightarrow n_{a0}^*$ and $n_r^* \rightarrow n_{r0}^*$ | $\text{s}^{-1}$ | $1.744 \times 10^{-1}$ <sup>a</sup> | LSR |
| $\gamma_{01}^*$ | Kinetic constant describing the rate of transitions $n_{a0}^* \rightarrow n_a^*$ and $n_{r0}^* \rightarrow n_r^*$ | $\text{s}^{-1}$ | $2.198 \times 10^{-2}$ <sup>a</sup> | LSR |
| $\gamma_{ar}^*$ | Kinetic constant describing the rate of transition $n_{a0}^* \rightarrow n_{r0}^*$ | $\text{s}^{-1}$ | $1.890 \times 10^{-5}$ | LSR |
| $\gamma_{ra}^*$ | Kinetic constant describing the rate of transition $n_{r0}^* \rightarrow n_{a0}^*$ | $\text{s}^{-1}$ | $5.529 \times 10^{-3}$ | LSR |
| $\gamma_{on,a}^*$ | Kinetic constant describing the rate of transition $n_{free}^* \rightarrow n_{a0}^*$ | $s^{-1}$ | $6.050 \times 10^{-5}$ | LSR |
| $\gamma_{on,r}^*$ | Kinetic constant describing the rate of transition $n_{free}^* \rightarrow n_{r0}^*$ | $s^{-1}$ | $6.816 \times 10^{-5}$ | LSR |
| $\gamma_{off,a}^*$ | Kinetic constant describing the rate of transition $n_{a0}^* \rightarrow n_{free}^*$ | $s^{-1}$ | $3.289 \times 10^{-7}$ | LSR |
| $\gamma_{off,r}^*$ | Kinetic constant describing the rate of transition $n_{r0}^* \rightarrow n_{free}^*$ | $s^{-1}$ | $4.201 \times 10^{-5}$ | LSR |
| $\gamma_{free \rightarrow st}^*$ | Kinetic constant describing the rate of transition $n_{free}^* \rightarrow n_{st}^*$ | $s^{-1}$ | $7.246 \times 10^{-3}$ | LSR |
| $\gamma_{st \rightarrow free}^*$ | Kinetic constant describing the rate of transition $n_{st}^* \rightarrow n_{free}^*$ | $s^{-1}$ | $1.817 \times 10^{-5}$ | LSR |
| $\gamma_{free \rightarrow dif}^*$ | Kinetic constant describing the rate of transition $n_{free}^* \rightarrow n_{dif}^*$ | $s^{-1}$ | $7.307 \times 10^{-3}$ | LSR |
| $\gamma_{dif \rightarrow free}^*$ | Kinetic constant describing the rate of transition $n_{dif}^* \rightarrow n_{free}^*$ | $s^{-1}$ | $4.359 \times 10^{-4}$ | LSR |
| $\gamma_{dif \rightarrow st}^*$ | Kinetic constant describing the rate of transition $n_{dif}^* \rightarrow n_{st}^*$ | $s^{-1}$ | $5.281 \times 10^{-3}$ | LSR |
| $\gamma_{st \rightarrow dif}^*$ | Kinetic constant describing the rate of transition $n_{st}^* \rightarrow n_{dif}^*$ | $s^{-1}$ | $2.240 \times 10^{-5}$ | LSR |
| $j_{tot,x=0}^b$ | Dimensionless total flux of tau from the soma into the axon | | $3.654 \times 10^{-3}$ | LSR |
| $n_{free,x=0}^c$ | Dimensionless concentration of free (cytosolic) tau at the axon hillock | | $6.030 \times 10^{-4}$ | LSR |
| $A$ | Coefficient in Eq. (S1) | | $3.254 \times 10^{-1}$ | LSR |

<sup>a</sup> Values of $\gamma_{10}^*$ and $\gamma_{01}^*$ given in Table S2 are effective values of these kinetic constants, which give the best fit of model predictions with experimental or interpolated data. The dwell time of tau on MTs may be as short as 40 ms (Janning et al. 2014; Igaev et al. 2014).

**Table S3.** Dimensionless dependent variables in the model of tau transport.

| Symbol | Definition |
| --- | --- |
| $j_{tot}$ | $\frac{j_{tot}^*}{n_{tot,x=0}^* v_a^*}$ |
| $n_{tot}$ | $\frac{n_{tot}^*}{n_{tot,x=0}^*}$ |
| $n_a$ | $\frac{n_a^*}{n_{tot,x=0}^*}$ |
| $n_r$ | $\frac{n_r^*}{n_{tot,x=0}^*}$ |
| $n_{a0}$ | $\frac{n_{a0}^*}{n_{tot,x=0}^*}$ |
| $n_{r0}$ | $\frac{n_{r0}^*}{n_{tot,x=0}^*}$ |
| $n_{free}$ | $\frac{n_{free}^*}{n_{tot,x=0}^*}$ |
| $n_{st}$ | $\frac{n_{st}^*}{n_{tot,x=0}^*}$ |
| $n_{dif}$ | $\frac{n_{dif}^*}{n_{tot,x=0}^*}$ |

### S2. Estimating the flux of tau into the axon terminal

To estimate the flux of tau into the axon terminal, 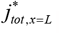, we used the approach developed in Li et al. (2012) for modeling the transport of neurofilament proteins. Tau that enters the axon terminal can either be degraded or reverse its direction of motion to retrograde and re-enter the axon. If 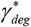 is a kinetic constant characterizing tau degradation rate and 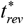 is the time required for a motor-driven tau to reverse its direction in the terminal, then the probability of tau degradation is estimated as 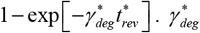 is estimated as 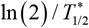 and 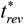 is estimated as 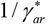. Since this estimate is approximate, we multiplied this estimate by a parameter *A*, as we did in Kuznetsov and Kuznetsov (2017a). The value of parameter *A* is determined by finding the best fit with published experimental data. This leads to the following equation:

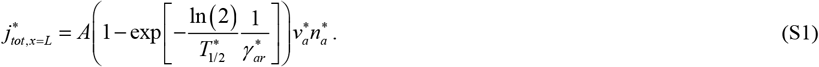

### S3. Numerical solution of equations simulating SAT of tau and finding the best-fit parameter values

Eqs. (1)-(7) compose a system that contains ordinary differential equations and algebraic equations. We used algebraic Eqs. (3), (4), and (7) to eliminate 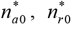, and 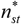 from the remaining equations. The remaining equations were solved subject to boundary conditions (12), (13) using the MATLAB’s BVP4C solver (MATLAB R2019a, MathWorks, Natick, MA, USA), Kuznetsov and Kuznetsov (2018).

The penalty function given by Eq. (14) was minimized using MultiStart with the local solver fmincon, which are part of MATLAB’s Optimization Toolbox. We used the default interior-point algorithm option in fmincon, which is described in Dantzig and Thapa (2003). 10,000 randomly selected starting points were used in MultiStart.

### S4. Simplified equations

#### S4.1. Simplified equations for 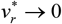

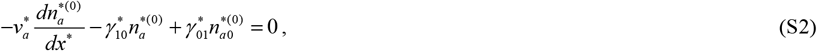

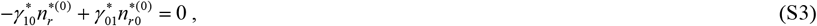

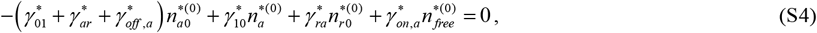

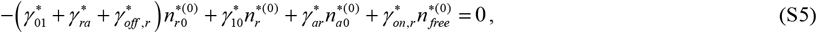

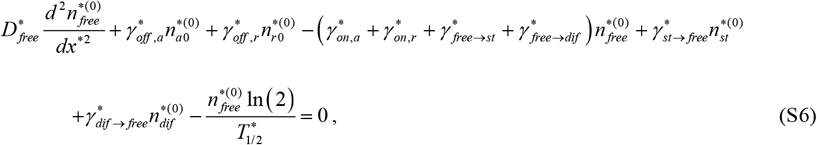

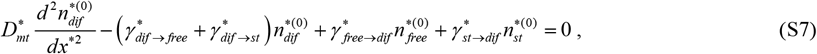

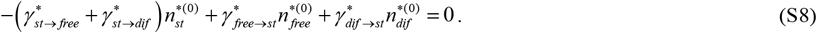

#### S4.2. Simplified equations for 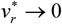 and 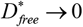

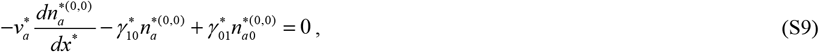

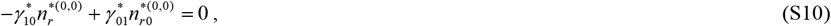

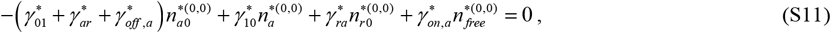

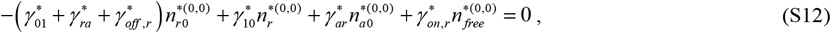

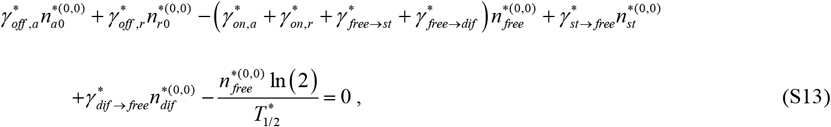

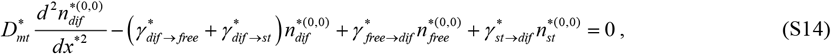

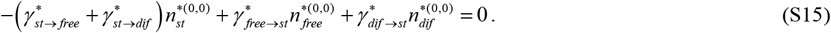

#### S4.3. Simplified equations for 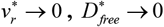, and 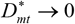

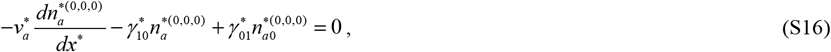

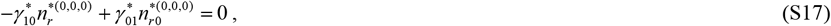

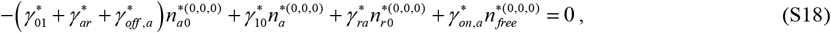

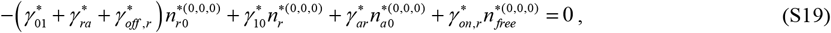

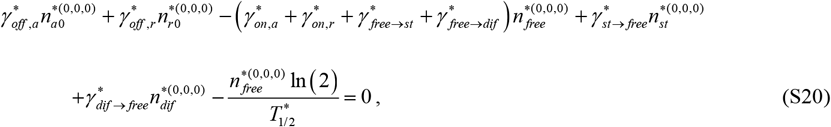

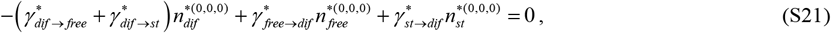

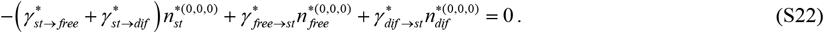

#### S4.4. Simplified equations after 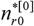 was eliminated using Eq. (S17)

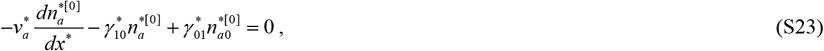

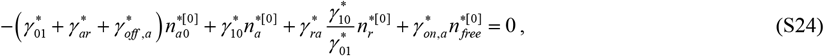

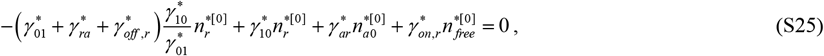

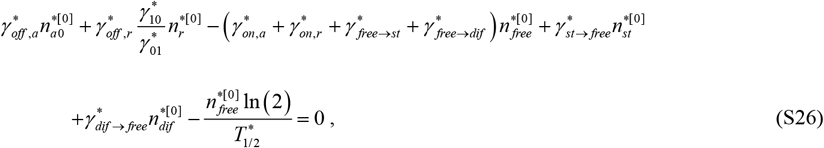

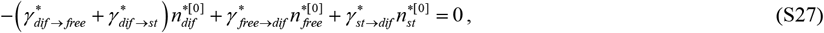

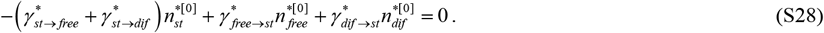

### S5. Supplementary figures

**Fig. S1.**
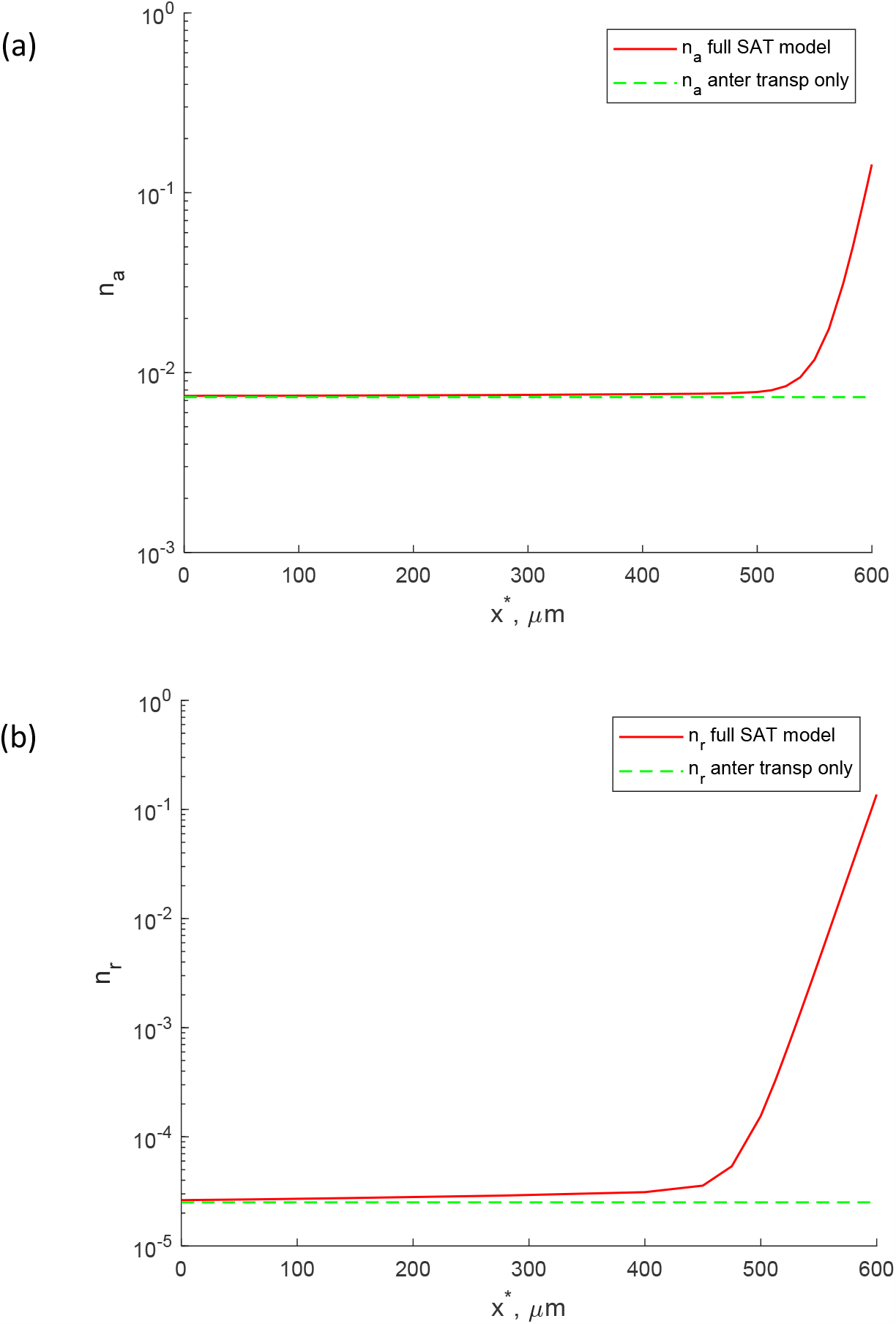
(a) Concentration of anterograde kinesin-driven tau. (b) Concentration of retrograde dynein-driven tau. Distributions for the full SAT model and the model that simulates anterograde motor-driven transport only are displayed.

**Fig. S2.**
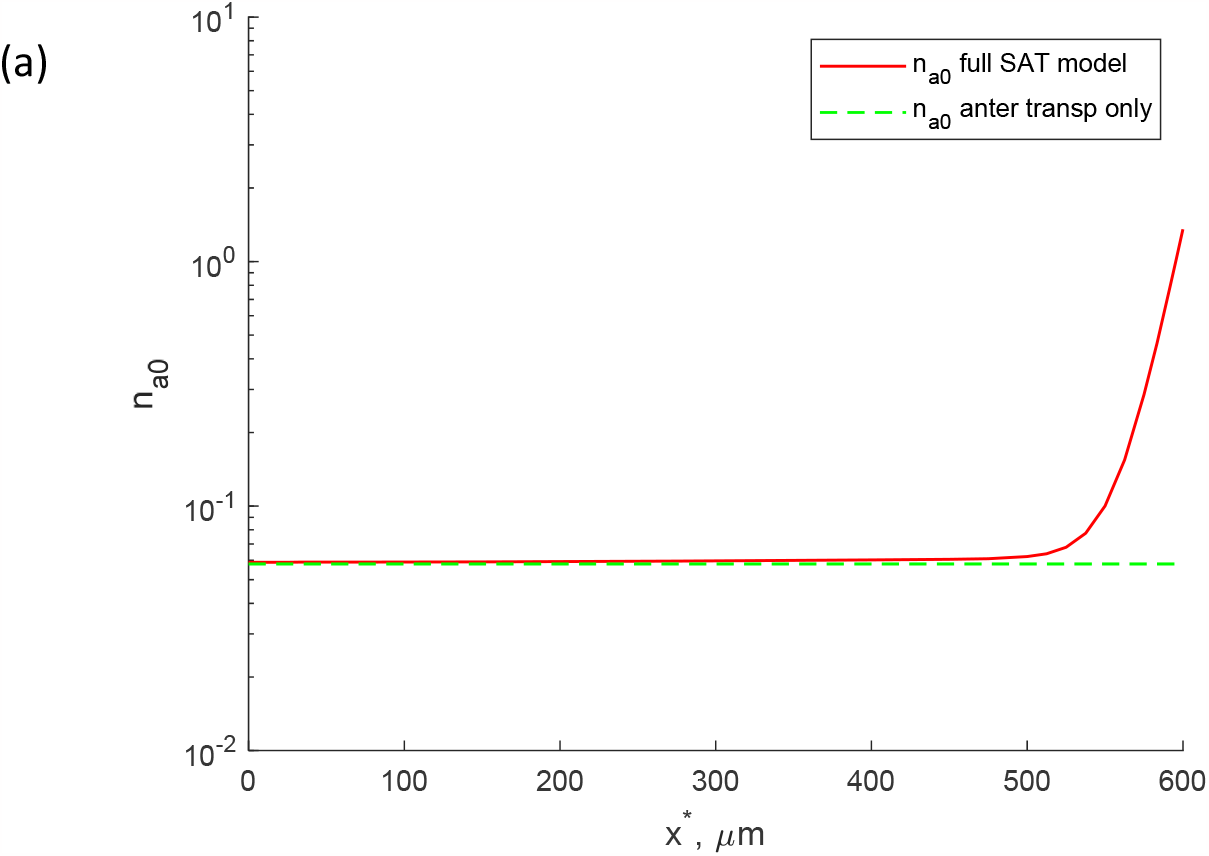
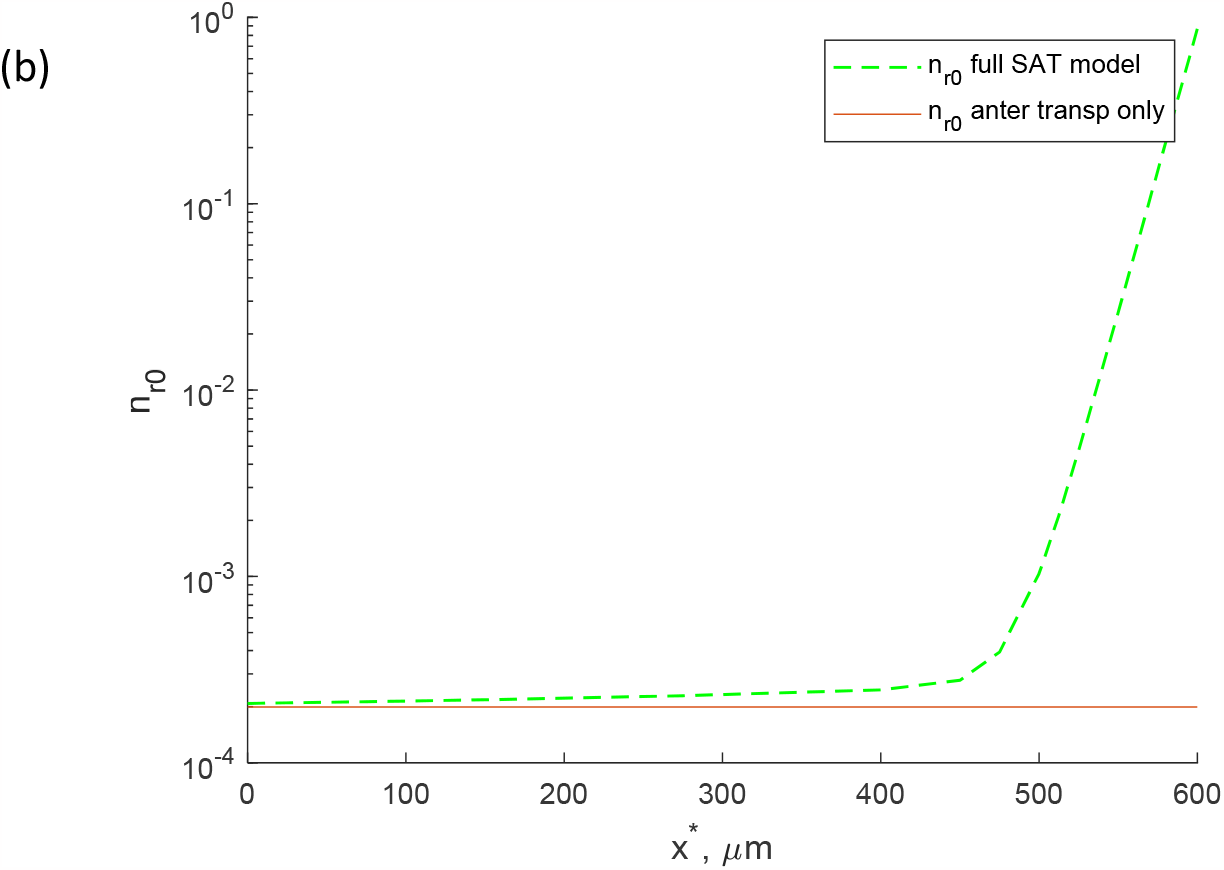
(a) Concentration of anterograde pausing tau. (b) Concentration of retrograde pausing tau. Distributions for the full SAT model and the model that simulates anterograde motor-driven transport only are displayed.

**Fig. S3.**
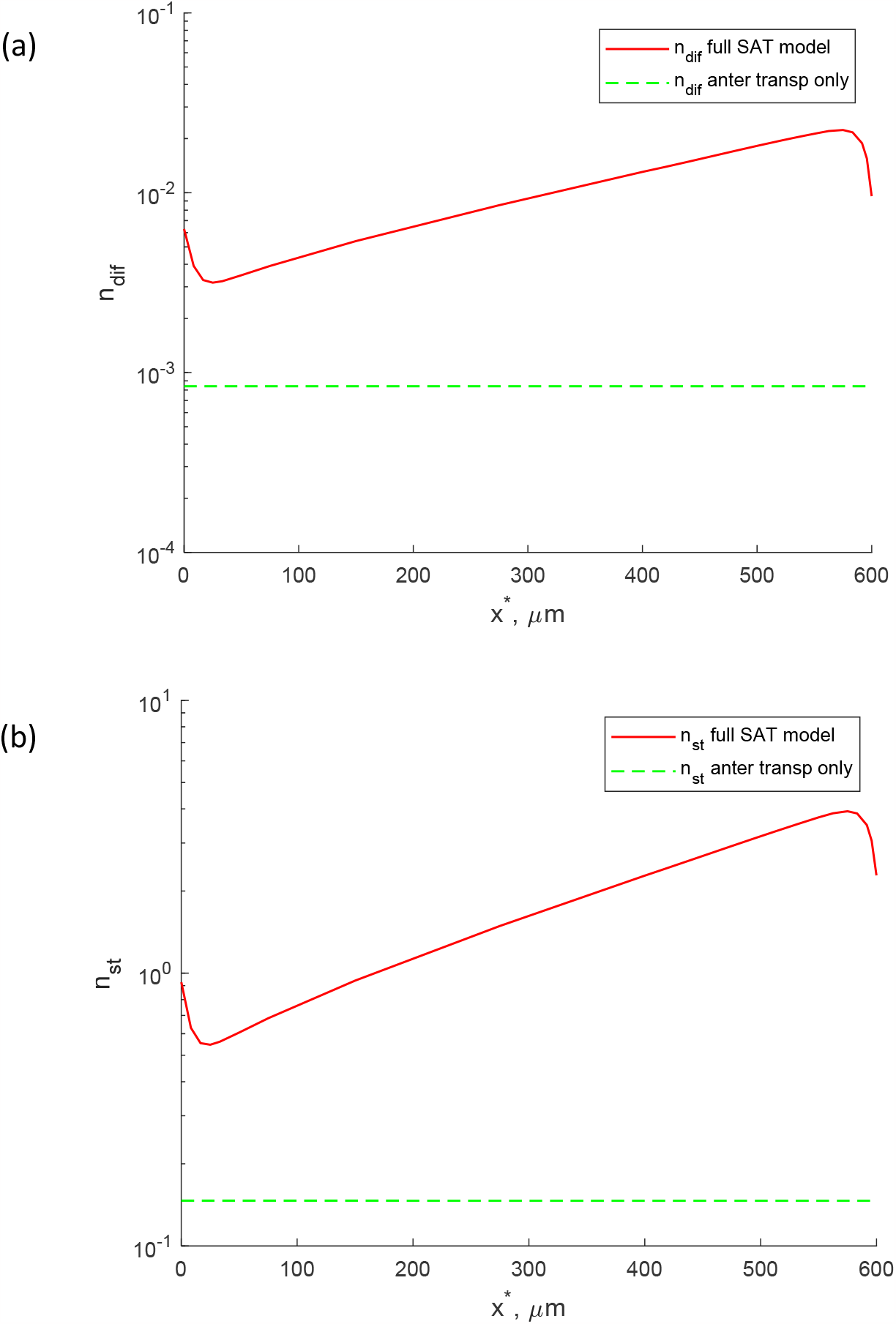
(a) Concentration of MT-bound tau that can diffuse along MTs. (a) Concentration of MT-bound tau that is stationary on MTs.

## Notes

### Competing Interest Statement

The authors have declared no competing interest.

### Summary of Updates

Clarified the novelty of our study, added an explanation about tau interaction with microtubules, and added some suggestions for future research.

